# Experimental drought reduces the productivity and stability of a recovering calcareous grassland

**DOI:** 10.1101/2023.07.11.548537

**Authors:** J. Jackson, S.L Middleton, C. S. Lawson, E. Jardine, N. Hawes, K. Maseyk, R Salguero-Gómez, A. Hector

## Abstract

1. Grasslands comprise 40% of terrestrial ecosystems and are globally important for food production, carbon storage, and other ecosystem services. However, grasslands in many areas are becoming increasingly exposed to extreme wet and dry periods resulting from global temperature increases.
2. Therefore, understanding how grasslands will respond to climate change is a pressing issue for managing changes to biodiversity and ecosystem service provision.
3. Here, we use experimental manipulations of precipitation (50% increase and 50% decrease of growing-season precipitation) to investigate the resistance of the diversity and productivity of a calcareous grassland community recovering from historical agricultural conversion.
4. We found that decreasing growing season precipitation led to reductions of mean productivity (25 % decrease in peak above-ground biomass) and its temporal stability (54 % increase in biomass variance across years). However, the grassland community composition was resistant to the precipitation manipulations, with no clear difference in community compositional turnover, dissimilarity, or biodiversity indices. Furthermore, the precipitation manipulations had no effect on the path of ongoing (30 year) recovery of grassland plant diversity from the period of previous agricultural conversion.
5. While the diversity of this calcareous grassland was resistant to precipitation extremes (at least in the short term), sustained reductions in growing-season precipitation reduced productivity and its temporal stability demonstrating that different properties of grasslands can vary in their responses to changes in precipitation.

## 1. Introduction

Over one million species are threatened with extinction due to rising global temperatures (Díaz et al., 2019). However, climate change is not universally associated with biodiversity loss (Antão et al., 2020). For plant communities, complex responses to climate change from the individual to the community level highlight the need for targeted community-level experiments (Gupta et al., 2020; Harrison et al., 2015; Kardol et al., 2010; Parmesan & Hanley, 2015). One of the key components of climate change that is expected to influence plant communities is more extreme precipitation patterns, and particularly drought (Hopkins & Prado, 2007; Knapp et al., 2015). For example, in grasslands precipitation is predicted to have a much larger impact than both temperature and CO_2_ increases, which are expected to have greater impacts on legume species (Hopkins & Prado, 2007). Therefore, there is a need to investigate how plant communities respond to changes in the climate, and in particular changing precipitation patterns (Franklin et al., 2016).

Grasslands are important ecosystems in which to explore responses to precipitation change. Approximately 40% of terrestrial ecosystems are grasslands (O’Mara, 2012; Petermann & Buzhdygan, 2021), and in Europe, grasslands occupy over 56 million ha (33%) of the agricultural area (European Commission, 2008). Furthermore, grasslands provide 38 ecosystem services, accounting for one third of carbon stored in terrestrial ecosystems, controlling soil erosion and mitigating against floods (Abberton et al., 2010; Bengtsson et al., 2019; Petermann & Buzhdygan, 2021; Zhao et al., 2020). However, grasslands have generally declined in extent, use, and biodiversity over the 20^th^ and 21^st^ centuries due to agricultural intensification, grazing, and conversion to other crops (Ceballos et al., 2010; Hilker et al., 2014; Peeters, 2009). The global importance of grasslands coupled with the threat of reduced stability with changes to precipitation (Fay et al., 2008) highlights the need to investigate how grasslands respond to a changing climate.

One striking example of a vulnerable grassland ecosystem is calcareous grasslands, the focus of this study. Calcareous grasslands are typified by basic soils, often occurring on limestone or chalk bedrock, supporting up to 700 vascular plant species in Europe, and providing a wide range of ecosystem services such as pollination, carbon sequestration, and recreation (Gibson & Brown, 1991; Grêt-Regamey et al., 2014; Klaus et al., 2021; Willems, 1990). There has been declines in the extent of many European calcareous grasslands, following agricultural intensification, scrub encroachment and grazer management (Grêt-Regamey et al., 2014; Ridding et al., 2020). To preserve high levels of species diversity, calcareous grasslands are listed as recovering ecosystems and part of wider conservation management schemes in countries such as the United Kingdom (Gibson, 1986; Gibson & Brown, 1991; Maddock, 2008; Poschlod et al., 1998).

Previous work has found evidence for both reductions in, and resistance of, grassland community diversity and productivity with changes in precipitation (and drought). A key study by Harrison et al. (2015) found that reductions in midwinter precipitation over 14 years reduced biodiversity in Californian grasslands. Furthermore, alongside work on grasslands, there has been a wide effort in plant community ecology to understand the influence of drought conditions on biomass/primary productivity, in particular using experimental manipulations of precipitation (Knapp et al., 2017; Kröel-Dulay et al., 2022). Generally, drought conditions reduce primary productivity in floral communities, including grasslands (Herben et al., 1995; Kardol et al., 2010; Kröel-Dulay et al., 2022; Wang et al., 2007). This reduction in primary productivity also impacts pollinator resources (Phillips et al., 2018), with expected impacts on community diversity. Yet, despite observed changes in floral biodiversity and productivity with drought in some grassland communities (Harrison et al., 2015), others have proved resistant to both experimentally-induced, and natural variation in drought or precipitation (Cleland et al., 2013; Craine et al., 2013; Grime et al., 2000, 2008). In particular, over a single year at the current study site, experimental removal of precipitation in July and August did not reduce species richness but instead decreased plant cover (with increases to litter cover) (Sternberg et al., 1999). Resistance to drought in native grasslands may be due to high species diversity conferring a range of physiological drought tolerance mechanisms, or increased presence of mycorrhizal fungi, buffering communities against the impacts of increased precipitation (Craine et al., 2013; Craven et al., 2018; Jia et al., 2021; Wagg et al., 2017).

Here, we use six-years (2016-2021) of biodiversity data from an experimental manipulation (50% increase and 50% decrease) of precipitation in a calcareous grassland recovering from disturbance to examine their resistance to this important component of climate change. Specifically, we test three key questions: i) whether changes in precipitation affect aboveground annual net primary productivity (aboveground ANPP) and its temporal stability across years, ii) whether calcareous grassland communities have a high resistance to drought conditions, and iii) whether changes in precipitation alter secondary successional pathways and community composition during calcareous grassland recovery from agricultural intensification. Furthermore, we investigate how changes in the plant communities were driven by site-level weather patterns, and changes in the abundance of individual species. We address these questions by quantifying community stability and resistance, which are a useful way to integrate community responses to climate change and compare communities from different ecosystems (Allen et al., 2019; Pimm, 1984; Van Meerbeek et al., 2021). Here, we quantify community stability by tracking changes in community composition and productivity through time with respect to environmental change (Donohue et al., 2013; Pimm, 1984). Importantly, shifts in the mean and variability of productivity or abundance (*e.g.* biomass) serve as early warning signals of community shifts, and indicate decreased stability in ecological communities (Clements & Ozgul, 2016; Pimm, 1984). We quantify stability at the community level by investigating how perturbations in precipitation influence compositional turnover (community change through time) and resistance (difference in community before and after a perturbation) (Donohue et al., 2013; Pimm, 1984).

## 2. Methods

### 2.1 Study site

The RainDrop (rainfall and drought platform) experiment is situated in a ∼2 ha area (‘five-acre field’) in the Upper Seeds grasslands (51°46’16.8”N 1°19’59.1”W, 166 m a.s.l) on the upper area of the Wytham hill in the University of Oxford’s Wytham estate, Oxfordshire, UK (Fig. S1). Upper Seeds is a recovering calcareous grassland, which was used for arable agriculture from the second world war until the late 1970s, before the site was managed as a grassland beginning in 1978 (Gibson, 1986; Grime et al., 2000). Upper Seeds, as with other calcareous grasslands, has a high level of floral biodiversity, in which graminoids constitute ∼60% of species by biomass. Management consists of mowing all above-ground vegetation in mid-July at the peak of the growing season, and again in early October, coinciding with the end of the growing season. Biomass is removed following mowing. The site has a shallow soil depth (300-500 mm), alkaline soils (Gibson & Brown, 1991), a daily average temperature range of −5 °C to 26 °C (2016-2020), and a daily total precipitation range of 0-40 mm (2016-2020) (Rennie et al., 2017).

### 2.2 Experimental design

We explored grassland biodiversity responses to precipitation in the context of the global drought network (DroughtNet) international drought experiment. DroughtNet’s international drought experiment is a coordinated distributed experiment with over 100 sites globally (https://drought-net.colostate.edu/). The goal of this experimental network is to explore ecosystem sensitivity to precipitation extremes through experimental manipulations of precipitation (Knapp et al., 2017). Precipitation manipulation is carried out by modifying natural precipitation patterns in each plot with rainout shelters, acting as a press disturbance (continuous change in environment) maintained across several years. The manipulation was implemented in RainDrop as a randomised, replicated block design in which four treatments were repeated across five blocks (n = 20 experimental plots; Fig. 1).

**Figure 1.**
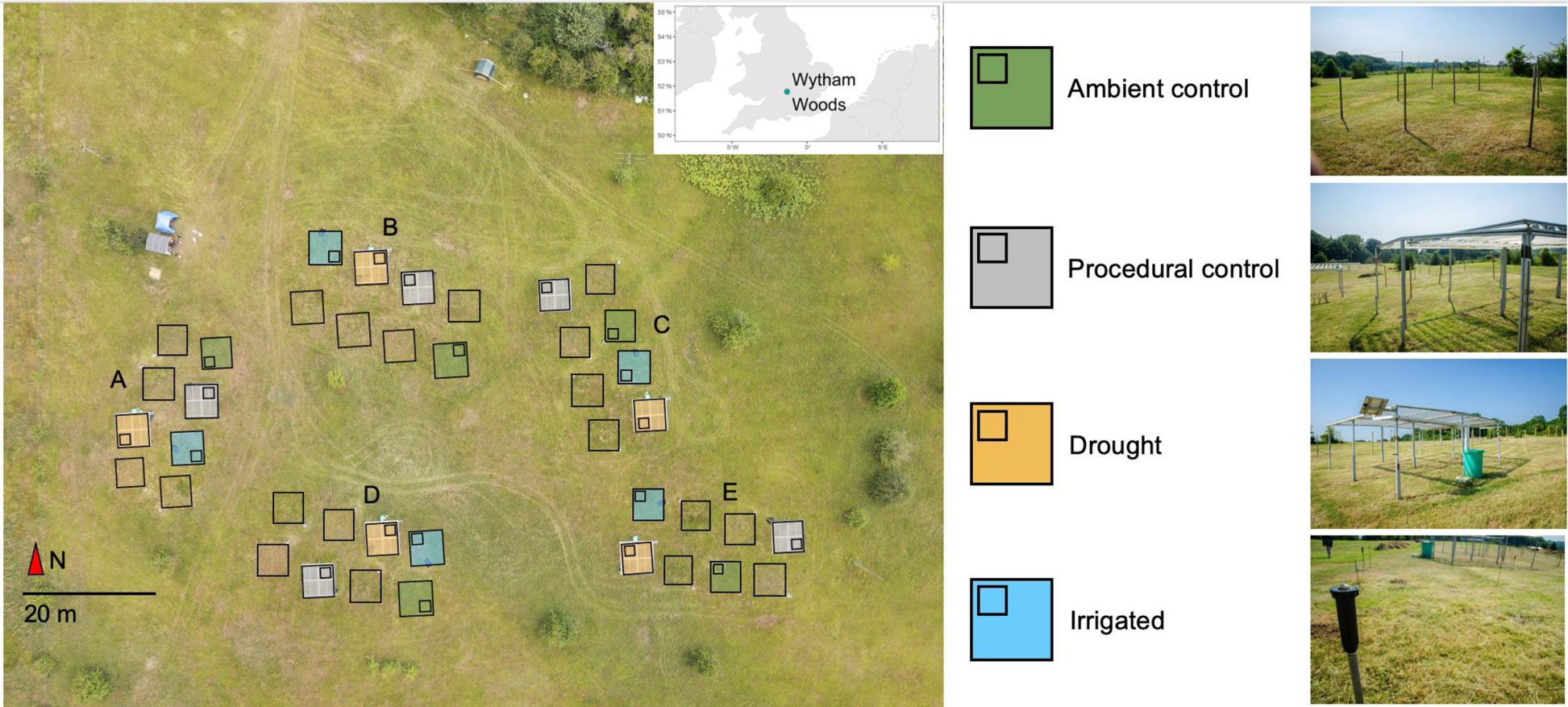
Experimental schematic of RainDrop on Upper Seeds, Wytham Woods with DroughtNet coordinated distributed experiment plots. DroughtNet treatments are denoted by the colour of each 5 m × 5 m plot, and include: Ambient control (green; no manipulation), Procedural control (grey; rainfall shelter but no change rainfall), Drought (orange; −50% rainfall shelter), and Irrigated (blue; +50% rainfall with sprinklers). Photos give a ground-level indication of each treatment type. Smaller squares indicate the biodiversity data collection quadrat in each plot, the positions of which were randomised. Additional transparent 5 m × 5 m plots indicate other experimental plots that are not included in the DroughtNet experiment.

Each replicated unit of 5 m × 5 m plots had one of four experimental treatments: ambient control plots (Ambient Control), −50% precipitation rainout shelters to simulate drought (Drought), +50% irrigated plots with sprinklers to simulate increased precipitation (Irrigated), and procedural controls (Procedural Control; precipitation shelter with no change to precipitation) (Fig. 1). Biodiversity data collection occurs in the central 1 m × 1 m quadrat in one quarter of each 5 m × 5 m plot, where the data collection quarter was randomised at the beginning of the experiment.

Rainout shelters consist of metal structures 1.5-2 m above the ground with transparent Perspex guttering. In the drought treatment, the rainout shelters are spaced such that 50% of the surface area of the plot is blocked by guttering. Irrigated treatment plots are supplied by water containers from the drought treatment, which collects precipitation with the transparent Perspex gutters, such that the 50% additional precipitation is comprised of precipitation lost to the shelter. Procedural Control plots test potential confounds of the drought treatment, in which similar structures and guttering are in place to simulate the micro-habitat of the shelter, but with guttering inverted to allow natural precipitation levels (Fig. 1). Precipitation manipulation percentages were selected through an assessment of long-term precipitation records, which found that extremes of annual precipitation differed from average years by ∼40% (Knapp et al., 2015). At RainDrop, all drought treatments are removed between October and March of each year, when the Perspex gutters are inverted to restore natural precipitation levels, such that the experimental treatments are active during the growing season. However, rainout shelters and guttering remain throughout the year.

### 2.3 Data collection

The core experimental protocol consists of biodiversity monitoring within experimental plots, namely, biomass (productivity) and species abundance. To explore how precipitation manipulation influences grassland dynamics and composition, we monitored three main features of biodiversity: total community above-ground annual net primary productivity (ANPP), functional group level aboveground biomass, and species-level percentage cover of vascular plants, for each 1 m × 1 m quadrat in each year between 2016-2021. We collected biomass at the peak of the growing season, 20^th^ June – 14^th^ July, and at the end of September, such that above-ground biomass measures give an annual estimate of above-ground net primary productivity across the growing season. We estimated ANPP using a ‘clip strip’ of all vascular plant material in a 1 m × 0.25 m strip in the centre of each quadrat, collected after percentage cover data. Clip strips were gathered using hand trimmers ∼ 1 cm above the soil surface. Within one day of collection, we sorted clip strips in to five functional groups: graminoids, legumes, non-leguminous forbs, woody species, and bryophytes and dried them at 70°C for 48 hrs, before weighing the dry biomass with an accuracy of ± 0.1g. Forbs are defined as herbaceous flowering plant other than a grass. We used both functional group-level estimates of biomass and summed values of ANPP in analyses, which were scaled by a factor of four to the standardised measure of g m^-2^. Due to smaller biomasses estimated for woody (1.65 % of total biomass) and bryophyte groups (1.20 % of total biomass), we only included graminoids, legumes and forbs in subsequent ANPP analyses.

Percentage cover data collection occurred at the peak of the growing season in each year, between 15^th^-30^th^ June. We estimated the percentage cover of all vascular plant species in each quadrat, and data from individual species was categorised as one of four broad functional groups: graminoids, legumes, forbs, and woody species. Because species overlapped spatially, percentage cover estimates exceed 100%. Species names were resolved using the International Plant Names Index (IPNI, 2022).

We added environmental context and explored how biodiversity changes are influenced by local weather patterns using weather data from the National Environment Research Council (NERC) Environmental Change Network (Rennie et al., 2017). A meteorological station was present in five-acre field within 100 m of all experimental quadrats. Raw meteorological data consisted of 16 weather variables, which were gathered at hourly intervals between 2016-2020, but data were not available in 2021. We used the mean hourly precipitation and temperature in the spring (21^st^ March – 20^th^ June) and summer (21^st^ June – 22^nd^ September) (the growing season) for each year of study as weather variables of interest.

### 2.4 General analysis

We analysed the experimental data with hierarchical Bayesian regression models using the *brms* package (Bürkner, 2017) in R version 4.1.3 (R Core Team, 2022). To perform model selection, we estimated the out-of-sample predictive performance of candidate models relative to base models that excluded predictor variables of interest. For each candidate model, we performed leave-one-out cross validation with the *loo* criterion and the expected log-wise predictive density (*elpd*, where Δ*elpd* gives the change in *elpd* relative to another explanatory model) (Vehtari et al., 2017). Therefore, *elpd* gives an estimate of predictive performance that is analogous to information criterion. Where two candidate models were comparable in *elpd* (Δ*elpd* < 2), we reported the model with fewer explanatory variables and explored the posterior coefficients of the model to make inference. Models were run across four Markov Chain Monte Carlo chains for 4,000 iterations with 2,000 warm-up iterations, and the convergence of the model across chains was assessed by inspecting *R* values, which assesses the degree of mixing between chains (Bürkner, 2017). For the full set of model priors, please refer to model code (10.5281/zenodo.8135588). Following model selection, we performed a set of Bayesian hypothesis tests (Bürkner, 2017) to investigate whether there were consistent differences in biodiversity measures between control treatments, and the proportion of variance explained by the random effect of experimental block. Differences between control treatments were evaluated by comparing posterior distributions between Ambient Control and Procedural Control groups. We used the intraclass correlation coefficients (ICC) (Nakagawa & Schielzeth, 2010) to assess the proportion of variance explained by block random effects relative to the total population-level variance.

### 2.5 Testing the effect of precipitation on grassland stability

To test our first key question, we analysed how precipitation manipulation influences ANPP and its stability. We examined responses at both the whole community level and at the functional group level, as well as the inter-annual temporal stability of above-ground ANPP. We define the temporal stability of productivity as the inverse of the interannual coefficient of variation (CV) of above-ground ANPP (He et al., 2022) for each quadrat. Where the response variable was the total and group-level ANPP, raw ANPP values were transformed using the natural log, which were modelled using a Gaussian distribution (although similar results were obtained using a Gamma distribution). In the ANPP models, the key predictors of interest were precipitation treatment and observation year. We used model selection to test the predictive performance for a set of candidate models including precipitation treatment (categorical variable, four levels), a linear effect of the observation year (continuous variable, z-scored), an autoregressive term for the observation year (order = 1), and two-way interactions between precipitation treatment and observation year (linear). We compared candidate models to base models that excluded predictor variables of precipitation treatment and observation year. ANPP models estimated at the level of functional group also included terms for functional group (categorical variable, three levels). The full set of candidate models for each ANPP response variable is detailed in supplementary tables S1-3. We also included a categorical predictor term for the month of harvest (middle or later part of growing season).

In all models, we included a clustered (hierarchical), intercept-only random effect of the precipitation treatment within block (five levels) to account for the experimental structure of RainDrop, and an intercept-only random effect of observation year (six levels) to capture additional interannual variability. For models of the temporal stability of ANPP, because the metric captured interannual variability in biomass for each quadrat, we could not include any temporal effects in model selection. Thus, for temporal stability models we tested a candidate model with the precipitation treatment to the base model with no predictor variables (Table S3). In ANPP models, we used weakly informed priors of *N*(3.5, 0.5) for the global intercept term and *N*(0, 1) for predictor variables. The intercept-only random effects were fitted using exponential priors with rates between 4-8.

### 2.6 Testing the resistance of grassland communities to precipitation treatments

We tested our second key question by investigating whether the grassland community was resistant to precipitation treatments in two ways: with broad diversity indices, and using community composition and turnover. First, we explored how broad diversity indices at the quadrat level were influenced by the precipitation treatments using linear hierarchical mixed-effects model selection, in an identical procedure as described in section 2.4. We calculated biodiversity indices using the relative proportions, *p*, of each species from percentage cover estimates. The three biodiversity indices included were vascular plant species richness, the Shannon-Weiner diversity index, *H* = − ∑ *p* ln *p* (Shannon & Weaver, 1963), and the Simpson’s diversity index, *D* = ∑ *p*^2^(Simpson, 1949). For the Shannon-Weiner and Simpson’s indices, response variables were z-scored (mean and variance centered on 0) for analyses and models were fitted using a Gaussian distribution. Models with species richness were fitted using a Poisson distribution and a log link. Shannon-Weiner and Simpson’s models were fitted with regularising priors of *N*(0, 0.5) for both intercept and predictor terms, and Richness models were fitted with a prior of *N*(3, 0.25) for the intercept.

To assess how the grassland community composition varied between precipitation treatments we tested community dissimilarity using non-linear multidimensional scaling (NMDS) implemented in the *vegan* package (Oskanen et al., 2022). Community-level data consisted of species percentage cover data for each quadrat observation, which is a single treatment in a given block each year. We fitted the NMDS using the Bray-Curtis dissimilarity index with a dimension of three, and up to 1000 random starts to reach convergence in stress values. Then, we compared the first two dominant NMDS axes between precipitation treatments. We used hierarchical linear mixed-effects models with a response variable of the NMDS axis scores, with model selection as described in section 2.4. We fitted these models with regularising priors of *N*(0,0.1) for both the global intercept and predictor variables. Then, in addition to the linear modelling framework, we explored statistical differences in Bray-Curtis similarities between precipitation treatments using analysis of similarities. Analysis of similarities tests differences in dissimilarity within sampling units compared to between sampling units (Oskanen et al., 2022). We also explored block-level community effects by testing the dissimilarity between experimental blocks.

### 2.7 Testing the recovery of the grassland community

We addressed our third key question by exploring whether there was evidence for recovery from agricultural management in the grassland community. We estimated temporal trends in diversity indices and NMDS axes, as well as the similarities in floral communities between observation years. We extracted temporal trends from linear models of biodiversity specified in section 2.5, which included linear or autoregressive effects of observation year, as well as two-way interactions between precipitation treatments and observation year. In addition to model selection, we also performed analysis of similarity tests for the NMDS communities between observation years (Oskanen et al., 2022). Therefore, by assessing the temporal change in diversity indices and NMDS scores, we tested the degree of recovery in this calcareous grassland.

### 2.8 Environmental and community context

Finally, to give further environmental and community-level context, we tested how local weather variables influenced ANPP and biodiversity indices, and which species were most important at driving community differences. Following the model selection framework, we implemented linear models to investigate the impact of mean temperature and precipitation across the spring and summer influenced biodiversity indices. In these models, linear terms for observation year were replaced with annual mean weather variables. We included both weather data for the current year and the previous year relative to biodiversity data collection, to test for current and lagged impacts of local weather on biodiversity differences.

To further explore the drivers of community composition differences between sampling groups of precipitation treatments and observation years, we investigated which species were most important for community differences. We calculated the Similarity Percentage for each species, which gives the decomposed contribution of individual species to the Bray-Curtis dissimilarity between communities (Oskanen et al., 2022). We tested Similarity Percentage for communities between precipitation treatment, observation year, and blocks. Then, following Similarity Percentage analysis, we explored variation in the relative abundance, *p*, of influential species between communities.

## 3. Results

### 3.1 Simulated drought reduces biomass production and its temporal stability

We found that simulated drought substantially reduces both ANPP and productivity’s temporal stability over time (Fig. 2). For total ANPP, the model with the highest predictive performance was the model including the categorical effect precipitation treatment (Δ*elpd* = 5.69 relative to the base model; Table S1). Total ANPP was substantially reduced in the drought treatment (*β* = −0.77 [−1.04; −0.49]; *β* coefficients give the posterior mean difference compared to the ambient control treatment on the log-scale, with the 95% credible intervals), with a mean total ANPP of 137 ± 156 (S.D.) g m^-2^ compared to 182 ± 137 (S.D.) g m^-2^ for the ambient control treatment (Fig. 2a). Thus, compared to ambient conditions, the mean ANPP was reduced by 24.7% in the drought treatment. There was weak negative skew in raw ANPP values, but this skew did not impact model convergence (Fig. S1). Furthermore, we re-ran model selection using a gamma distribution, which did not change the qualitative results (Fig. S1b). We observed a similar pattern for temporal stability in productivity, with a Δ*elpd* of 3.15 for the model including precipitation treatment compared to the base model (Table S3). Temporal stability in productivity was substantially reduced in the drought treatment (*β* = −0.19 [−0.32; −0.06]), equating to a 53.5% increase in the coefficient of variation of biomass (CV = 0.56) relative to the ambient treatment (CV = 0.37) (Fig. 2b). There were no discernible differences in temporal stability in productivity in either the control treatment or irrigation treatment.

**Figure 2.**
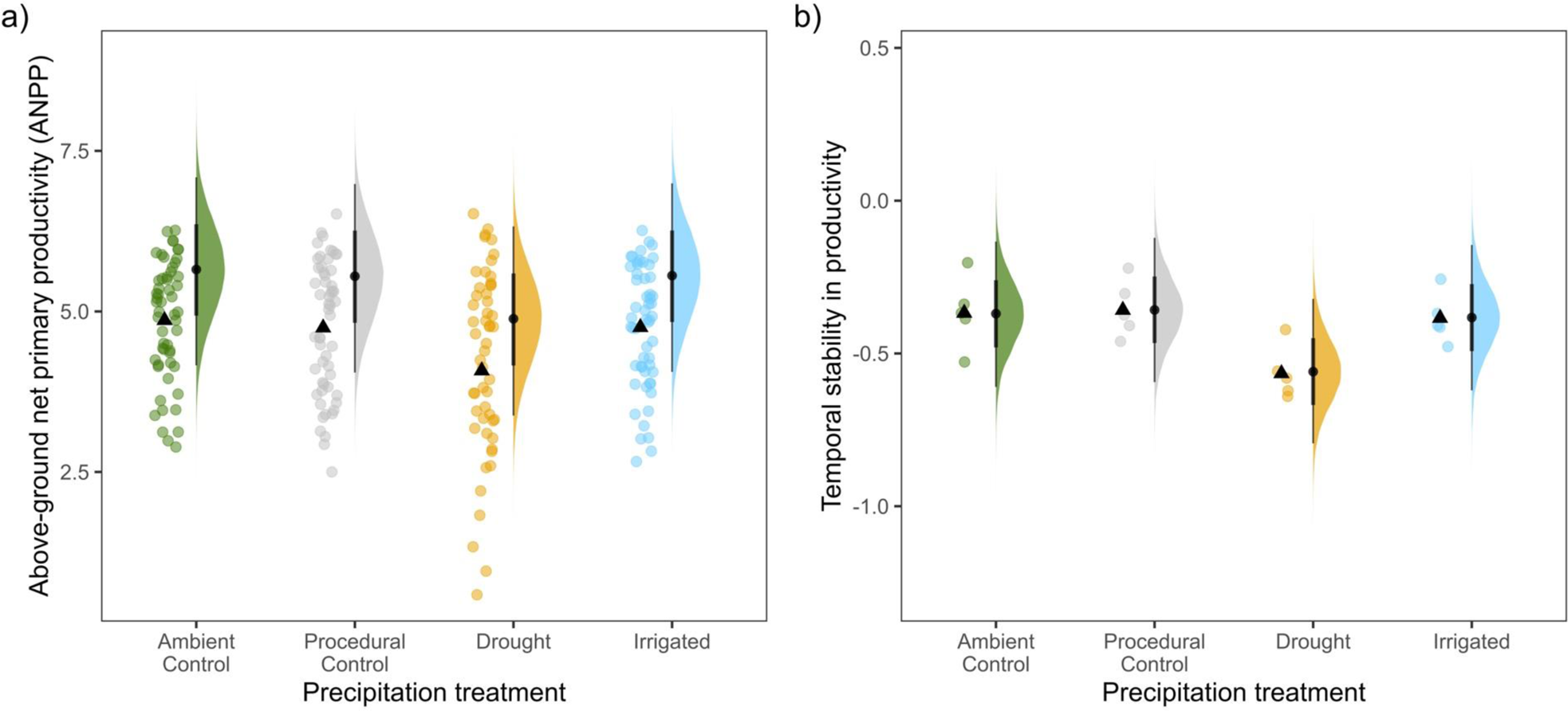
Drought reduces average ANPP and its temporal stability. A) Total above-ground annual net primary productivity (aboveground ANPP) with respect to precipitation treatments, where ANPP is the natural log-transformed above-ground biomass in g m^-2^. B) Temporal stability in productivity (inverse of interannual coefficient of variation in biomass) with respect to precipitation treatment. Coloured points give raw data across blocks and years and black triangles give the mean total ANPP. There is negative skew in raw ANPP data, which leads to reductions in mean ANPP values relative to predictions, but this skew did not impact model convergence or influence findings (Fig. S2). Distributions are derived from 8000 draws of the full posterior distribution including random effects, with probability density function boxplots giving the posterior mean and uncertainty.

The reductions in Overall ANPP were driven primarily by decreases in the biomass of graminoids and legumes (Table S2; Fig. S2). Forbs did not exhibit ANPP reductions in the drought treatment (*β* = −0.03 [−0.46; 0.39]), but reductions were accentuated in both graminoids (*β* = −1.00 [−1.41; −0.59]) and forbs (*β* = −0.87 [−1.29; −0.46]) (Fig. S2). Therefore, we observed mean reductions in ANPP of 36.1% and 36.4% for graminoids and legumes, respectively. Both mean total ANPP and group-level ANPP were overlapping between ambient and procedural control treatments (*β* = 0.11 [−0.17; 0.38], *β* = 0.31 [−0.11; 0.72]; differences in posterior means between ambient and procedural control). Furthermore, we did not find substantial variance in total biomass between blocks (ICC or *σ*_*block*_ = 0.01 [0.00; 0.05]). Finally, we did not observe any clear patterns between average spring and summer weather conditions and ANPP (Fig. S3).

### 3.2 Resistance of community diversity and composition to drought and irrigation

We found evidence that species composition of these calcareous grassland communities are resistant to precipitation treatments, both in terms of broad diversity indices and community composition (Fig. 3). For species richness, the Shannon-Weiner index, and the Simpson’s index, we did not observe differences in indices between precipitation treatments (Fig. 3a). For all three indices, the Δ*elpd* compared to the base model was below 0.6, indicating no clear association between the indices and precipitation treatments, so we retained models excluding precipitation treatment (Table S4-6). Similarly to ANPP, there were no clear differences in ambient and procedural controls for richness, Shannon-Weiner index, or Simpson’s index. Furthermore, the mean posterior block-level variance (ICC or *σ*_*block*_) was below 0.03 for all indices. Similarly to ANPP, average spring and summer weather conditions were not strongly associated with any of the biodiversity indices (Fig. S4).

**Figure 3.**
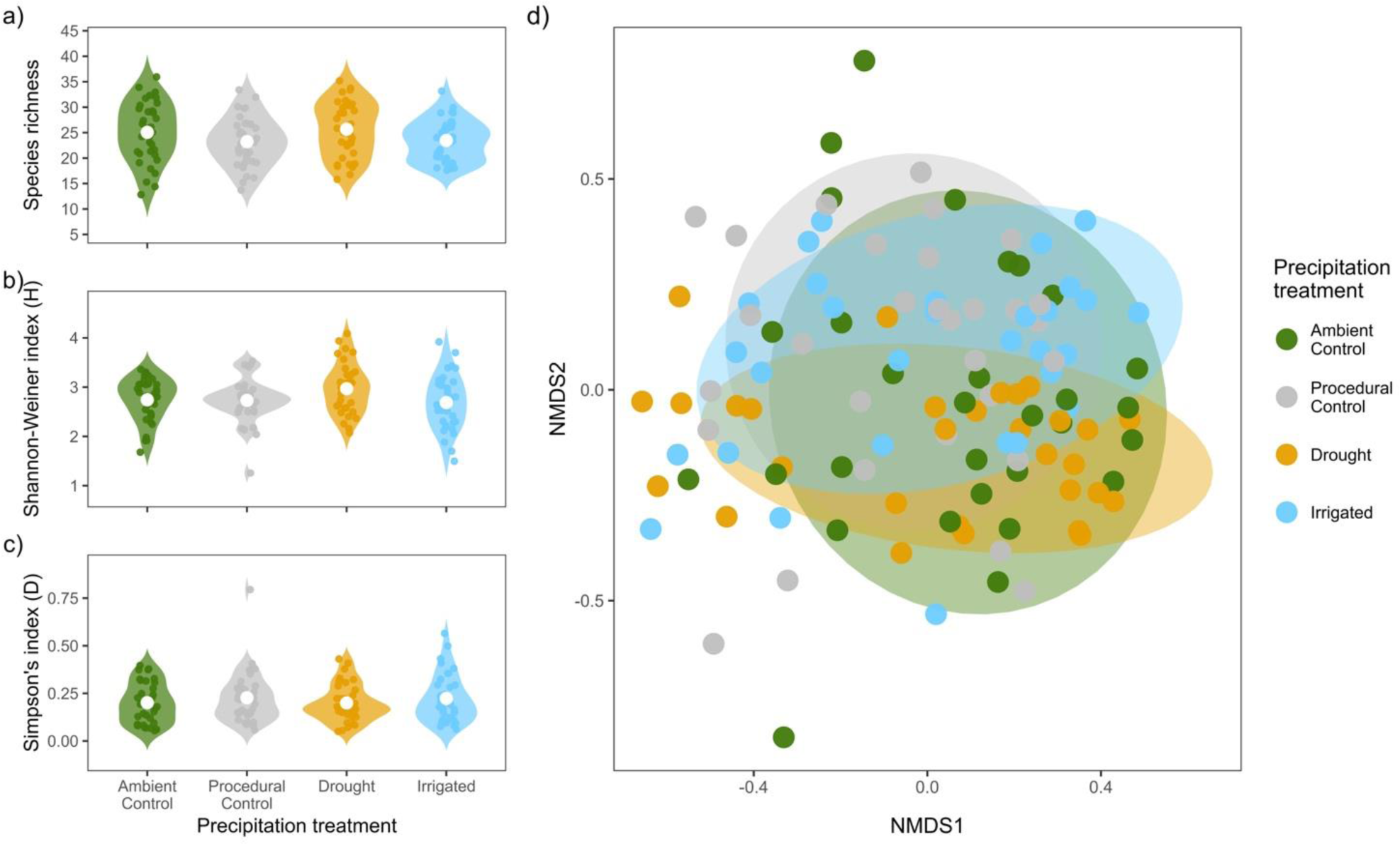
Calcareous grassland community diversity is resistant to precipitation manipulations. A-c) raw data distributions for species richness (a), Shannon-Weiner index (b) and Simpson’s index (c) with respect to precipitation treatment. Coloured points indicate raw data, and violins give an estimate of data density across each index. White points indicate mean biodiversity index values. d) Non-metric multi-dimensional scaling (NMDS) results for community composition, where the first two axes (NMDS1 & NMDS2) are displayed with respect to precipitation treatments. Ellipses are the 80% two-dimensional quantiles of the NMDS axes.

Community composition was also not clearly associated with the precipitation treatments. There was no clear association between NMDS axes one and two (NMDS1 and NMDS2) and precipitation treatment (Fig. 4a; Fig. S5), or between precipitation treatments and NMDS3 (Fig. S5). This was further supported by the analysis of similarities, for which within sample dissimilarity was broadly comparable to dissimilarity between samples (Fig. S6; marginally significant relationship).

**Figure 4.**
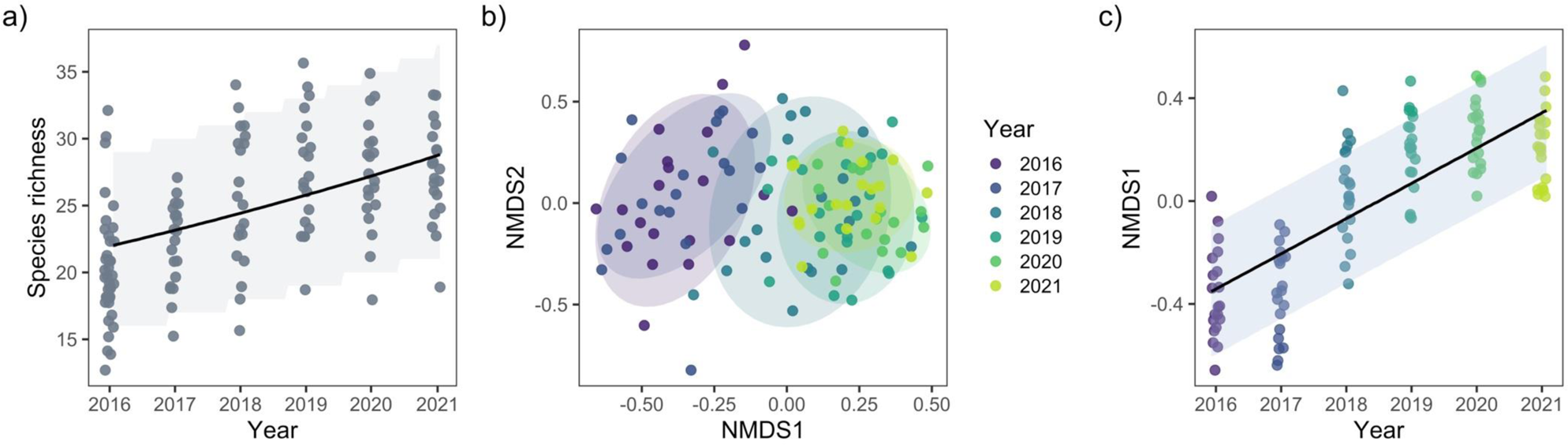
Changes in the grassland diversity and community composition 2016-2021. A) Increases in species richness between 2016-2021. Grey points give raw observations of species richness for each quadrat in each treatment. Solid line indicates the posterior mean prediction with 90% credible intervals, which include random effects. B) Non-metric multi-dimensional scaling (NMDS) results for community composition, where the first two axes (NMDS1 & NMDS2) are displayed with respect to observation year. Ellipses are the 80% two-dimensional quantiles of the NMDS axes. C) Increases to NMDS1 over the observation period. Points give observed NMDS1 scores for each quadrat for a given treatment and block across the study period. Solid black line indicates the posterior mean, with the 90% credible interval.

### 3.3 Recovery of the grassland from previous agricultural use

Across all observation years and experimental quadrats, there was a mean total above-ground net primary productivity (ANPP) of 303 ± 138 (S.D.) g m^-2^. Graminoids were the dominant functional group by biomass, with 63.6% of all biomass measured, compared to 20.4% for legumes and 16.0% for forbs. We recorded a total of 110 vascular plant species between 2016 and 2021 across all plots. At the quadrat level, species richness ranged between 13-36 species m^-2^ with a mean of 24.4 ± 5.14 (S.D.) species.

We found evidence that the calcareous grassland communities were undergoing secondary successional recovery from previous agricultural use between 1950-1978, with increases in species richness and changes in community composition (Fig. 4). However, these changes were not affected by the precipitation treatments. We found a positive association between species richness and observation year, such that there were increases in richness over the study period (Fig. 4a; *β* = 0.10 [0.04; 0.16]). Overall, between 2016 and 2021, there was an increase in mean richness from 20.3, to 27.4 species (Fig. 4a). Furthermore, we found a strong association between community composition and observation year (Fig. 4b). The analysis of similarities indicated a large difference in within-year community differences compared to between year community differences (Fig. S6). Furthermore, we observed a strong positive association between NMDS1 and observation year, which was a substantially better predictive model compared to the base model (Δ*elpd* = 58.4). There was a consistent increase in NMDS1 over the study period (*β* = 0.23 [0.20; 0.27]), indicating a shift in community composition (Fig. 4c).

### 3.4 Key species for community dissimilarity

Finally, we explored similarity percentages across species to investigate the species that were most influential in driving differences in community dissimilarity between years. Ten species had mean percentage contributions to dissimilarity above 2%, of which four were graminoids, and five were forbs including four legumes (Fig. 5a). Of these species however, *Arrhenatherum elatius* (graminoid), *Brachypodium pinnatum* (graminoid) and *Lotus corniculatus* (legume) had mean contributions of over 5% (median > 6.5%) to community dissimilarity across years (Fig. 5a & b). These three species varied substantially across the study period, and in particular *Arrhenatherum elatius*, which varied between an average abundance of 13.3% per quadrat in 2018 to 2.3% in 2021 (Fig. 5b).

**Figure 5.**
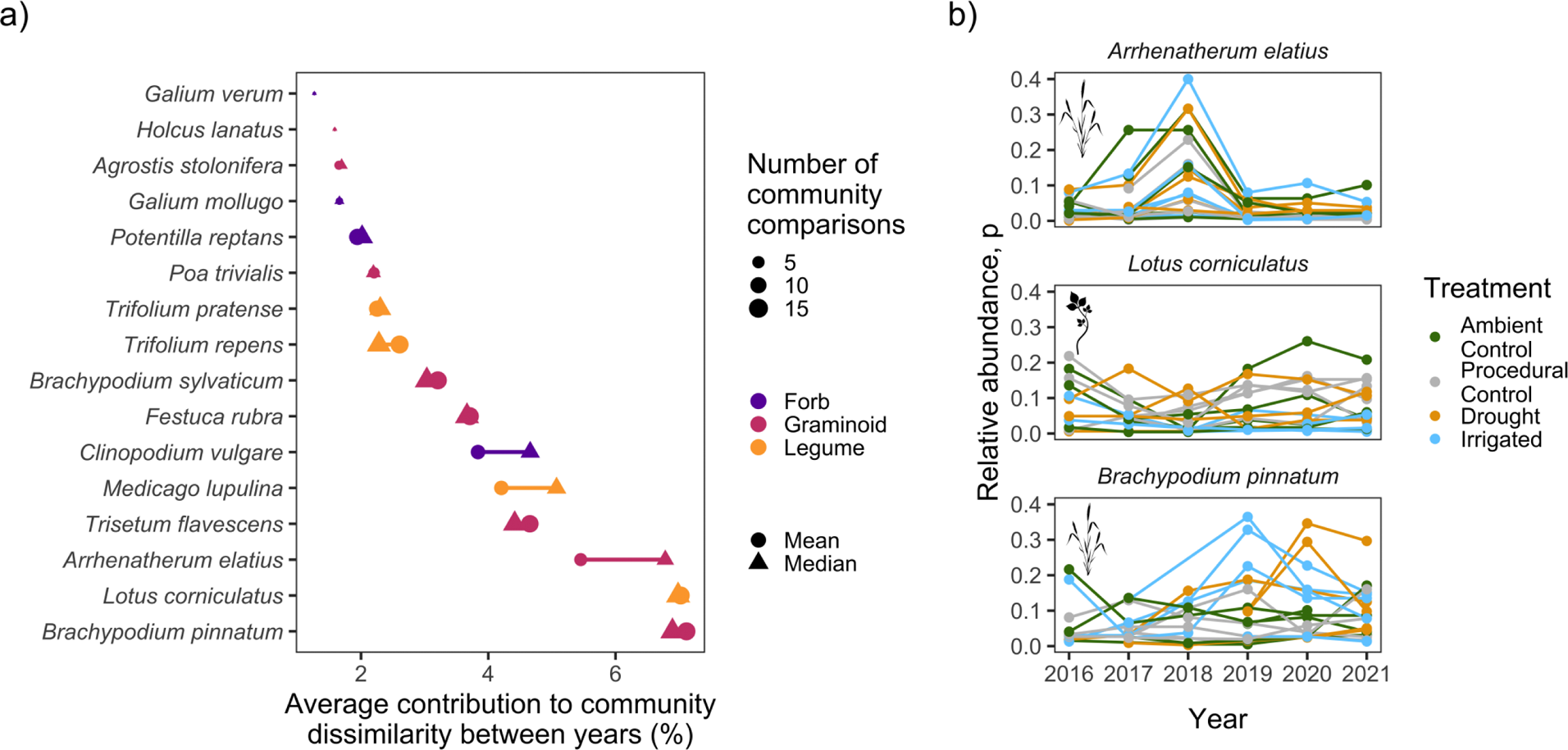
Commonly occurring graminoid and legume species drive community dissimilarity. A) Similarity percentage results for species driving community dissimilarity across years. Only 16 species (of 110) with the highest contributions are shown. Circles give mean percent contribution, and triangles mean percent contribution, with the size of the point indicating the number of community comparisons and the colour denoting the functional group. B) Temporal trends in the relative abundance of the three most influential species. Points and lines give raw estimates of relative abundance for each quadrat.

## 4. Discussion

From our six-year study of precipitation manipulation in a calcareous grassland, we show that simulated drought reduced the productivity and temporal stability of calcareous grasslands, but that species diversity and composition were generally resistant. These results add to global findings of decreased grassland productivity with drought (Kröel-Dulay et al., 2022), but also emphasise that the reductions in productivity are coupled with reduced temporal stability. While we don’t explicitly link biodiversity levels to the response of the grassland community to drought, our finding of community resistance is consistent with native British grasslands conferring a diversity of drought tolerance (Craine et al., 2013). Furthermore, our findings suggest that through reduced productivity, an increased prevalence of drought may impact key ecosystem services of calcareous grasslands, including pollination and carbon sequestration (Grêt-Regamey et al., 2014).

We found that drought conditions reduced the productivity of this calcareous grassland habitat and its inter-annual temporal stability, with 24% reductions in above-ground net primary productivity, and a 53% increase in productivity variance. Reductions of primary productivity and biomass in general have been widely reported in plant communities globally (Haddad et al., 2002; Kröel-Dulay et al., 2022). However, Kröel-Dulay et al. (2022) found that experimental manipulations of precipitation often underestimate the impact of drought on biomass, failing to capture other abiotic processes that are associated with drought in real-world settings. Increasing droughts threaten the future productivity of grasslands, which may be driven by altered patterns of nutrient cycling following precipitation extremes (Haddad et al., 2002). In addition to changes in mean productivity, drought also decreased the temporal stability by increasing the variance of productivity. Importantly, measures of stability can provide comparable metrics that indicate broader community-level changes as the environment changes (Clements & Ozgul, 2016; Wu et al., 2020). For example, He et al. (2022) found that experimental drought treatments in an alpine grassland had reduced stability during an extreme precipitation event. Studies explicitly linking climate change, grassland stability and ecosystem services will provide comparable insights into the future of plant communities globally.

Although we found evidence for reduced stability in productivity, over six years the diversity and composition of communities in this calcareous grassland were generally resistant to drought. Namely, we did not find any effect of precipitation extremes on community composition, turnover, or broad diversity indices that combined species richness and evenness. These findings are supported by findings from other grasslands, in which diversity and composition also proved resistant (Craine et al., 2013). Generally, native biodiverse ecosystems such as calcareous grasslands are predicted to have a high diversity of physiological drought tolerance, increasing community resistance to precipitation extremes through traits such as root thickness (Craine et al., 2013; Tucker et al., 2011). In an Alpine grassland, graminoids were more resistant to both drought and temperature manipulations than legumes (Tello-García et al., 2020). Drought resistance was also observed with 13 years of experimental manipulations of an unproductive, low diversity, grazed UK grassland (Grime et al., 2008). However, in contrast, 14 years of biodiversity data from Californian grasslands revealed close links between precipitation extremes and biodiversity (Harrison et al., 2015). The reason for differences in drought resistance between different grassland ecosystems is less clear, but resistance in calcareous grasslands may be the result of high biodiversity (a maximum of 51 species per 5 m^2^ compared to 33 per 5 m^2^ in Harrison et al. (2015)) if a wide range of ecological traits ensures resistance of the system as a whole (Hector et al., 2010). Furthermore, heterogeneity in soil is also linked to grassland resistance (Fridley et al., 2011). Our findings do not exclude broader level impacts from climate change on grassland communities, which may occur abruptly when thermal tolerance limits are reached (Trisos et al., 2020) over longer timescales, or in synergy with other drivers such as habitat fragmentation (Brook et al., 2008; Klaus et al., 2021).

In our study, richness increased by over 7 species m^-2^ between 2016-2021 indicating that these communities are still undergoing secondary successional recovery from previous conversion to agriculture that lasted from the 1950s until approximately 1978 (Gibson, 1986). However, while it is currently too early to ascertain non-linear dynamics in the change of community structure, the current data are consistent with a change in system state after 2019, leading to a plateau in community change. Future data collection will disentangle temporal dynamics in the community. Nevertheless, the precipitation manipulations could potentially alter the path of secondary succession but to date we find no evidence that the reductions in productivity have any associated effects on the on-going increases in species diversity and changes in species composition. Another potential explanation for the increase in richness is recovery from grazing, which was carried out intermittently on the site until more recent experiments (Gibson & Brown, 1991; Grime et al., 2000).

## Conclusions

Results from the first six years of precipitation manipulation demonstrate clear negative impacts of reducing precipitation by half on aboveground annual net primary production (ANPP) and its inter-annual temporal stability. This reduction in ANPP came about mainly due to decreases in the biomass of graminoids and legumes. However, to date the species diversity and composition of our grassland communities have proved resistant to reduced precipitation. On-going monitoring will be required to identify any longer-term consequences of the decrease in productivity caused by the reduction in precipitation.

## Acknowledgements

Special thanks to N. Fisher, N. Havercroft, and K. Crawford for field logistic support at Wytham throughout the study. Thanks also to M. Stone and D. Gowing for their work setting up and supporting the experiment. Also, thanks for assistance in the field from J. Haynes, L. Clements, L. McManus, H. King, D. Encarnation, A. Patwary, and L. Hinchcliffe. JJ was funded by the Amazon Web Service Test Bed Funding scheme “Monitoring and Predicting Biodiversity Resilience through AI & Robotics” to RS-G and NH and by a John Fell Funds grant to RS-G. RS-G was funded by a NERC IRF (NE/M018458/1). AH was supported by the John Fell Fund. AH, KM, and the Raindrop project were supported by the John Fell Fund, the Ecological Continuity Trust, the Patsy Wood Trust and the British Ecological Society.

## Declaration of Competing Interest

We declare that there are no competing interests.

## Data Availability Statement

All code, output and analysis data used in the current study are archived using the Zenodo repository DOI: 10.5281/zenodo.8135588, which was created from the following GitHub repository: https://github.com/jjackson-eco/raindrop_biodiversity_analysis. Raw biodiversity data (which is part of a global network) is available on request from Andrew Hector.

## Notes

https://zenodo.org/record/8135588

### Competing Interest Statement

The authors have declared no competing interest.

### Summary of Updates

Author typo and minor update

