## Supplementary Information for "Experimental drought reduces the productivity and stability of a recovering calcareous grassland"

Supplementary Information Contents:


| | | |
|---|---|---|
| Figure S1 | Exploring skew in ANPP | 2 |
| Figure S2 | Primary productivity separated by functional group | 3 |
| Table S1 | Model selection for ANPP | 4 |
| Table S2 | Model selection for group-level ANPP | 5 |
| Table S3 | Model selection for temporal stability of productivity | 5 |
| Table S4 | Model selection for the Shannon-Weiner index | 5 |
| Table S5 | Model selection for the Simpson’s index | 5 |
| Table S6 | Model selection for species richness | 6 |
| Table S7 | Model selection for NMDS axis 1 | 6 |
| Table S8 | Model selection for NMDS axis 2 | 6 |
| Table S9 | Model selection for NMDS axis 3 | 7 |
| Figure S3 | Weather effects on ANPP | 7 |
| Figure S4 | Weather effects on biodiversity | 8 |
| Figure S5 | NMDS precipitation treatment effects | 9 |
| Figure S6 | Analysis of community similarities | 10 |

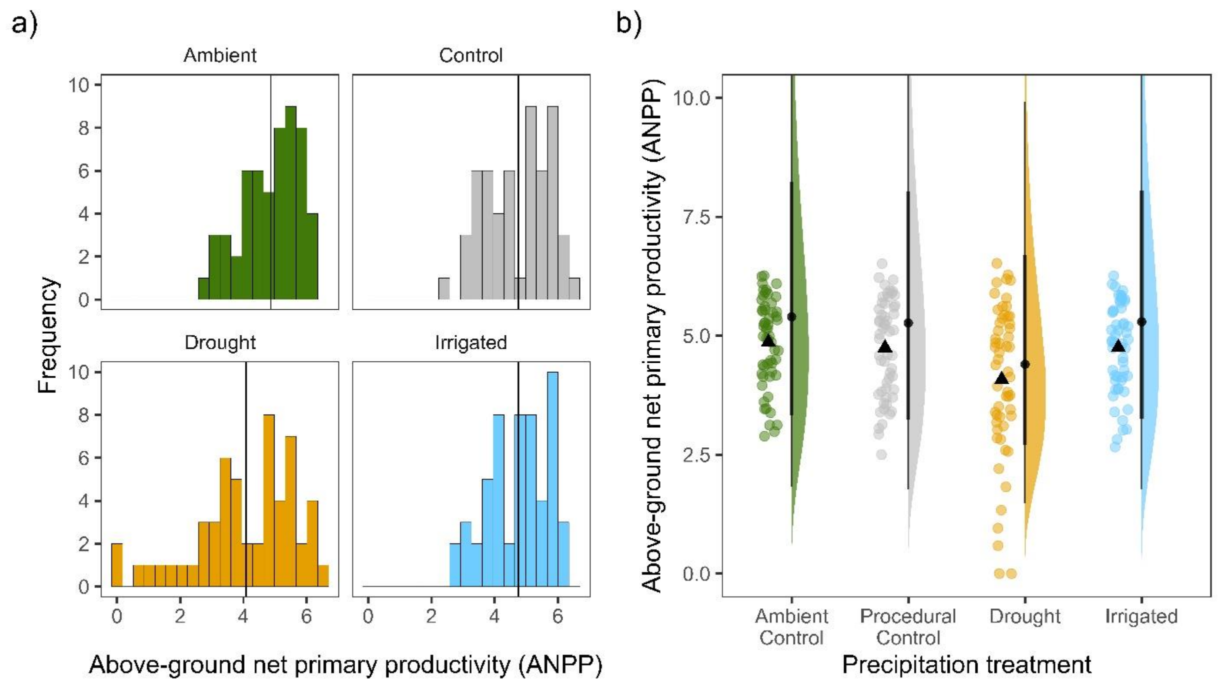


**Figure S1. Exploring skew in above-ground annual net primary productivity (ANPP).** ANPP is the natural log-transformed above-ground biomass in g $m^{-2}$. a) Frequency histograms for ANPP values in each precipitation treatment, indicating weak negative skew towards lower ANPP values. Based on this skew, we re-ran models of ANPP differences between precipitation treatments with gamma distribution models (shape = 5, scale = 0.8). b) Posterior predictions from gamma models of ANPP differences between precipitation treatments. Coloured points give raw data across blocks and years and black triangles give the mean total ANPP. Distributions are derived from 8000 draws of the full posterior distribution including random effects, with probability density function boxplots giving the posterior mean and uncertainty.

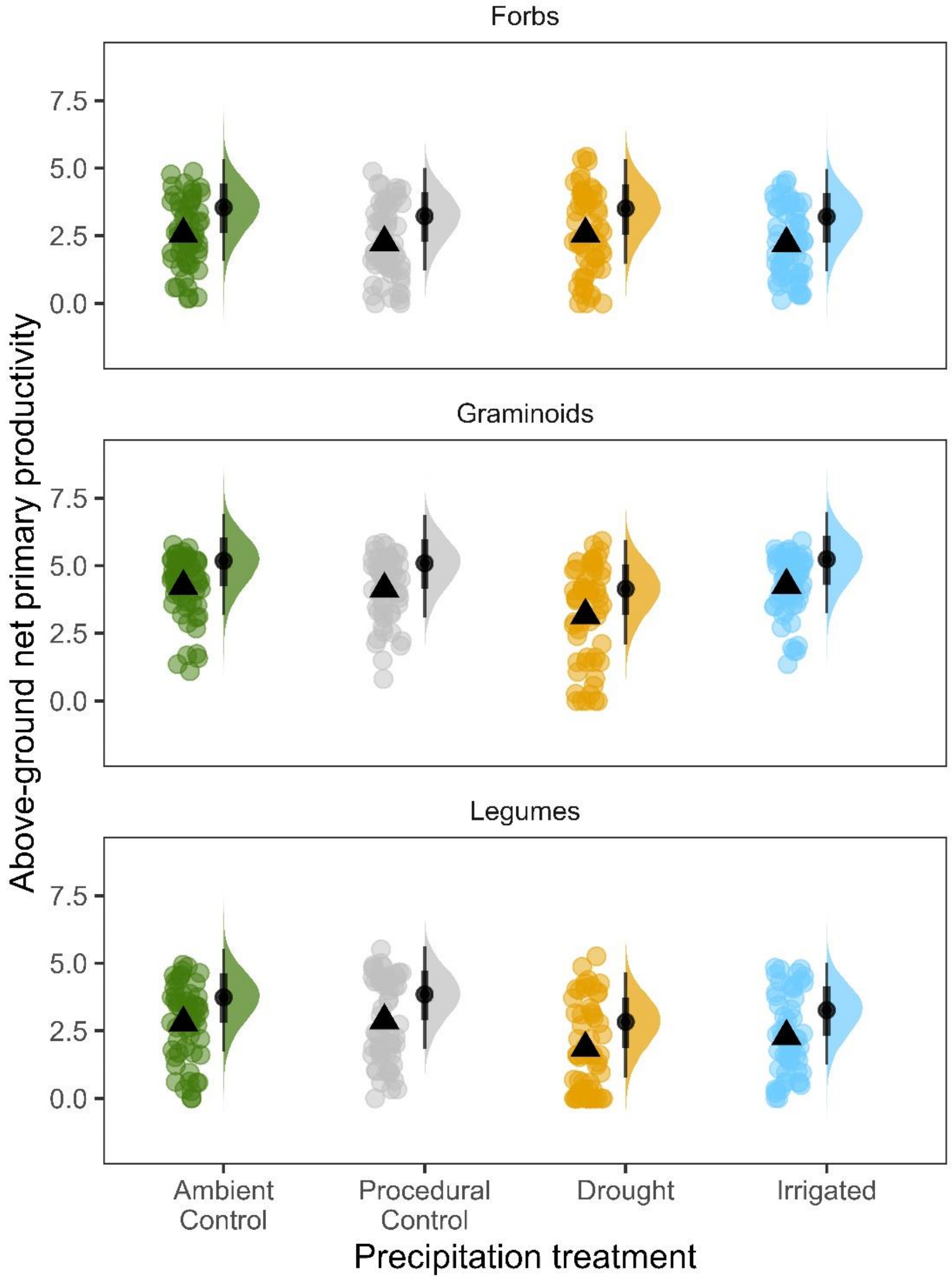


**Figure S2. Stability reductions are driven by Graminoids and legumes.** Differences in above-ground net primary productivity (ANPP) separated by functional group, between precipitation treatments. ANPP is the natural log-transformed above-ground biomass in g $m^{-2}$. Coloured points give raw data across blocks and years and black triangles give the mean total ANPP. Distributions are derived from 8000 draws of the posterior distribution, with probability density function boxplots giving the posterior mean and uncertainty.

Table S1. Model selection results for models of the total above ground annual primary productivity (ANPP; natural log of total biomass) total with respect to precipitation treatments and temporal trend effects. Models were compared using leave-one-out cross (LOO) validation and the expected log-wise predictive density ($elpd$). Column 'elpd difference' gives $\Delta elpd$, which was used as the measure of a model's predictive performance relative to the base model. Model formula is presented in hierarchical model syntax from the *lme4* package. *year_s* model term refers to a scaled continuous linear effect of observation year, and *year_f* refers to a categorical effect of year as a factor. The *ar()* term gives AR(1) autoregressive effects for observation year.

| R object | Model formula | LOO elpd | LOO elpd error | elpd difference | elpd error difference | LOO information criterion |
|---|---|---|---|---|---|---|
| totbiomass_year_linear | log_tot_biomass ~ 1 + treatment + harvest + year_s + (1 \| block/treatment) + (1 \| year_f) | -232.07 | 19.55 | 0.00 | 0.00 | 464.15 |
| totbiomass_treatment | log_tot_biomass ~ 1 + treatment + harvest + (1 \| block/treatment) + (1 \| year_f) | -232.31 | 19.65 | -0.24 | 0.21 | 464.62 |
| totbiomass_year_harvest | log_tot_biomass ~ 1 + treatment + year_s * harvest + (1 \| block/treatment) + (1 \| year_f) | -232.92 | 19.45 | -0.85 | 0.38 | 465.84 |
| totbiomass_year_treat | log_tot_biomass ~ 1 + treatment * year_s + harvest + (1 \| block/treatment) + (1 \| year_f) | -233.05 | 19.44 | -0.97 | 1.03 | 466.10 |
| totbiomass_base | log_tot_biomass ~ 1 + harvest + (1 \| block/treatment) + (1 \| year_f) | -237.34 | 21.85 | -5.27 | 4.04 | 474.68 |
| totbiomass_year_auto | log_tot_biomass ~ 1 + treatment + ar(gr = year_s, p = 1) + harvest + (1 \| block/treatment) | -239.43 | 17.69 | -7.36 | 5.87 | 478.86 |

Table S2. Model selection results for models of the group-level above-ground biomass with precipitation treatments and temporal trend effects.

| R object | Model formula | LOO elpd | LOO elpd error | elpd difference | elpd error difference | LOO information criterion |
|---|---|---|---|---|---|---|
| biomass_treatment_group_int | biomass_log ~ 1 + treatment * group + harvest + (1 \| block/treatment) + (1 \| year_f) | -816.57 | 20.54 | 0.00 | 0.00 | 1,633.14 |
| biomass_treatment_group_year | biomass_log ~ 1 + treatment * group * year_s + harvest + (1 \| block/treatment) + (1 \| year_f) | -817.71 | 20.12 | -1.14 | 3.82 | 1,635.42 |
| biomass_treatment_group | biomass_log ~ 1 + treatment + group + harvest + (1 \| block/treatment) + (1 \| year_f) | -845.65 | 21.76 | -29.08 | 7.71 | 1,691.30 |
| biomass_group | biomass_log ~ 1 + group + harvest + (1 \| block/treatment) + (1 \| year_f) | -846.27 | 21.87 | -29.70 | 7.96 | 1,692.55 |
| biomass_treatment | biomass_log ~ 1 + treatment + harvest + (1 \| block/treatment) + (1 \| year_f) | -1,026.00 | 17.12 | -209.43 | 15.40 | 2,052.01 |
| biomass_base | biomass_log ~ 1 + harvest + (1 \| block/treatment) + (1 \| year_f) | -1,026.75 | 16.90 | -210.18 | 15.49 | 2,053.49 |

Table S3. Model selection results for models of the temporal stability of productivity with respect to precipitation treatments.

| R object | Model formula | LOO elpd | LOO elpd error | elpd difference | elpd error difference | LOO information criterion |
|---|---|---|---|---|---|---|
| biomass_var_treat | cv_biomass ~ 1 + treatment + (1 ǀ block) | 14.43 | 2.72 | 0.00 | 0.00 | -28.86 |
| biomass_var_base | cv_biomass ~ 1 + (1 ǀ block) | 10.90 | 2.72 | -3.53 | 2.87 | -21.81 |

Table S4. Model selection results for models of the Shannon-Weiner index with respect to precipitation treatments and temporal trend effects.

| R object | Model formula | LOO elpd | LOO elpd error | elpd difference | elpd error difference | LOO information criterion |
|---|---|---|---|---|---|---|
| shannon_base | shannon ~ 1 + harvest + (1 ǀ block/treatment) + (1 ǀ year_f) | -188.54 | 9.54 | 0.00 | 0.00 | 377.09 |
| shannon_treatment_year_int | shannon ~ 1 + harvest + treatment * year_s + (1 ǀ block/treatment) + (1 ǀ year_f) | -189.77 | 9.17 | -1.23 | 2.41 | 379.54 |
| shannon_treatment_year_auto | shannon ~ 1 + harvest + treatment + ar(gr = year_s, p = 1) + (1 ǀ block/treatment) + (1 ǀ year_f) | -190.26 | 9.23 | -1.71 | 3.00 | 380.51 |
| shannon_treatment | shannon ~ 1 + harvest + treatment + (1 ǀ block/treatment) + (1 ǀ year_f) | -190.57 | 9.44 | -2.03 | 1.09 | 381.14 |
| shannon_year_linear | shannon ~ 1 + harvest + treatment + year_s + (1 ǀ block/treatment) + (1 ǀ year_f) | -190.69 | 9.38 | -2.15 | 1.09 | 381.39 |

Table S5. Model selection results for models of the Simpson's index with respect to precipitation treatments and temporal trend effects.

| R object | Model formula | LOO elpd | LOO elpd error | elpd difference | elpd error difference | LOO information criterion |
|---|---|---|---|---|---|---|
| simpsons_treatment_year_int | simpsons ~ 1 + harvest + treatment * year_s + (1 ǀ block/treatment) + (1 ǀ year_f) | -192.29 | 11.94 | 0.00 | 0.00 | 384.58 |
| simpsons_base | simpsons ~ 1 + harvest + (1 ǀ block/treatment) + (1 ǀ year_f) | -192.74 | 13.08 | -0.45 | 3.23 | 385.48 |
| simpsons_treatment_year_auto | simpsons ~ 1 + harvest + treatment + ar(gr = year_s, p = 1) + (1 ǀ block/treatment) + (1 ǀ year_f) | -192.90 | 12.45 | -0.61 | 3.91 | 385.80 |
| simpsons_treatment | simpsons ~ 1 + harvest + treatment + (1 ǀ block/treatment) + (1 ǀ year_f) | -193.10 | 12.60 | -0.81 | 2.61 | 386.19 |
| simpsons_year_linear | simpsons ~ 1 + harvest + treatment + year_s + (1 ǀ block/treatment) + (1 ǀ year_f) | -193.25 | 12.55 | -0.96 | 2.53 | 386.50 |

Table S6. Model selection results for models of species richness with respect precipitation treatments and temporal trend effects.

| R object | Model formula | LOO elpd | LOO elpd error | elpd difference | elpd error difference | LOO information criterion |
|---|---|---|---|---|---|---|
| richness_year_linear | richness ~ 1 + harvest + treatment + year_s + (1 l block/treatment) + (1 l year_f) | -399.62 | 5.29 | 0.00 | 0.00 | 799.25 |
| richness_base | richness ~ 1 + harvest + (1 l block/treatment) + (1 l year_f) | -399.96 | 5.20 | -0.33 | 1.52 | 799.91 |
| richness_treatment | richness ~ 1 + harvest + treatment + (1 l block/treatment) + (1 l year_f) | -400.40 | 5.23 | -0.78 | 0.96 | 800.81 |
| richness_treatment_year_auto | richness ~ 1 + harvest + treatment + ar(gr = year_s, p = 1) + (1 l block/treatment) + (1 l year_f) | -400.86 | 5.16 | -1.24 | 0.97 | 801.72 |
| richness_treatment_year_int | richness ~ 1 + harvest + treatment * year_s + (1 l block/treatment) + (1 l year_f) | -402.20 | 5.42 | -2.57 | 0.35 | 804.39 |

Table S7. Model selection results for models of community structure, captured with the first axis of non-linear multidimensional scaling (NMDS), with respect to temporal trend effects. The *block_treatment* effect indicates a categorical effect for the specific plot (a treatment within a block).

| R object | Model formula | LOO elpd | LOO elpd error | elpd difference | elpd error difference | LOO information criterion |
|---|---|---|---|---|---|---|
| nmds1_year | MDS1 ~ 1 + year_s + (1 l block/treatment) | 26.71 | 6.84 | 0.00 | 0.00 | -53.41 |
| nmds1_year_rs | MDS1 ~ 1 + year_s + (1 + year_s l block/treatment) | 26.30 | 7.02 | -0.40 | 1.67 | -52.61 |
| nmds1_year_blocktreatment | MDS1 ~ 1 + year_s * block_treatment + (1 l block/treatment) | 23.32 | 6.73 | -3.39 | 3.09 | -46.64 |
| nmds1_base | MDS1 ~ 1 + (1 l block/treatment) | -31.51 | 5.57 | -58.21 | 6.51 | 63.01 |

Table S8. Model selection results for models of community structure, captured with the second axis of non-linear multidimensional scaling (NMDS), with respect to temporal trend effects.

| R object | Model formula | LOO elpd | LOO elpd error | elpd difference | elpd error difference | LOO information criterion |
|---|---|---|---|---|---|---|
| nmds2_base | MDS2 ~ 1 + (1 l block/treatment) | 20.51 | 12.22 | 0.00 | 0.00 | -41.02 |
| nmds2_yearf | MDS2 ~ 1 + year_f + (1 l block/treatment) | 19.31 | 13.72 | -1.20 | 2.74 | -38.63 |
| nmds2_year | MDS2 ~ 1 + year_s + (1 l block/treatment) | 19.28 | 12.54 | -1.23 | 0.73 | -38.56 |
| nmds2_year_blocktreatment | MDS2 ~ 1 + year_s * block_treatment + (1 l block/treatment) | 18.52 | 11.10 | -1.99 | 4.14 | -37.05 |

Table S9. Model selection results for models of community structure, captured with the third axis of non-linear multidimensional scaling (NMDS), with respect to temporal trend effects.

| R object | Model formula | LOO elpd | LOO elpd error | elpd difference | elpd error difference | LOO information criterion |
|---|---|---|---|---|---|---|
| nmds3_yearf | MDS3 ~ 1 + year_f + (1 l block/treatment) | 20.23 | 8.03 | 0.00 | 0.00 | -40.45 |
| nmds3_year | MDS3 ~ 1 + year_s + (1 l block/treatment) | -5.41 | 7.20 | -25.64 | 5.48 | 10.82 |
| nmds3_base | MDS3 ~ 1 + (1 l block/treatment) | -5.71 | 6.72 | -25.93 | 5.41 | 11.41 |
| nmds3_year_blocktreatment | MDS3 ~ 1 + year_s * block_treatment + (1 l block/treatment) | -14.49 | 8.08 | -34.71 | 6.58 | 28.97 |

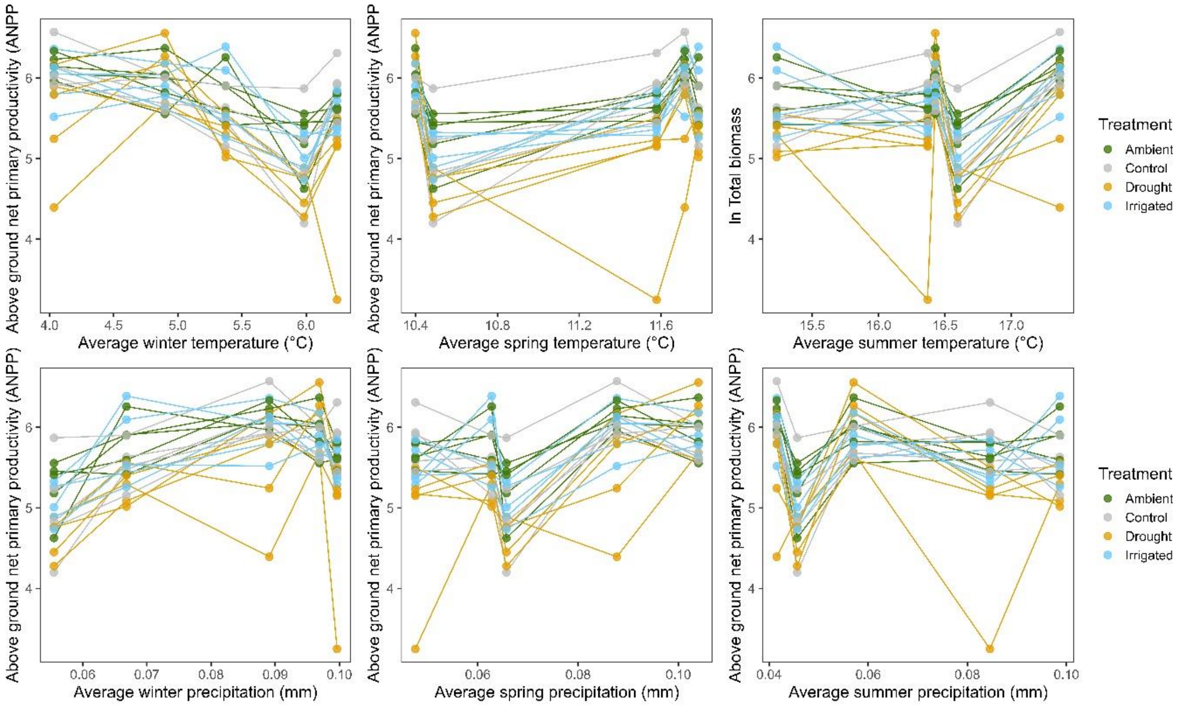


**Figure S3.** No clear effect of average daily precipitation and temperature in the winter, spring and summer on above-ground annual net primary productivity.

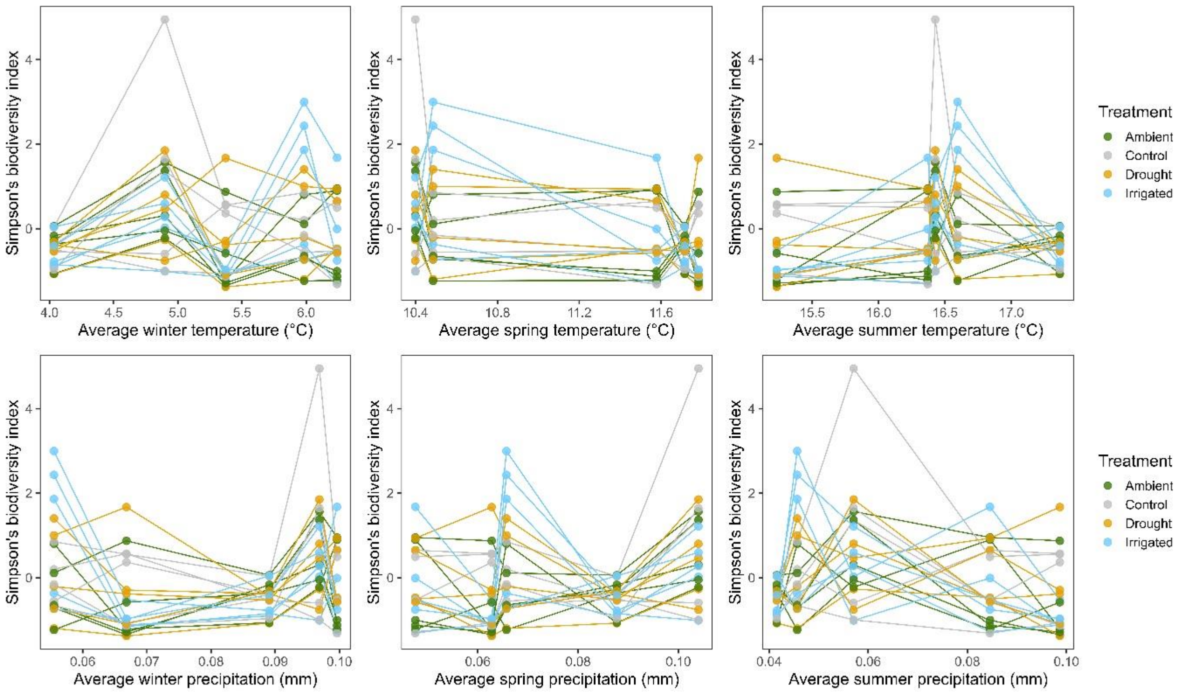


**Figure S4.** No clear effect of average daily precipitation and temperature in the winter, spring and summer on Simpson's biodiversity index. Similar patterns observed for other biodiversity metrics.

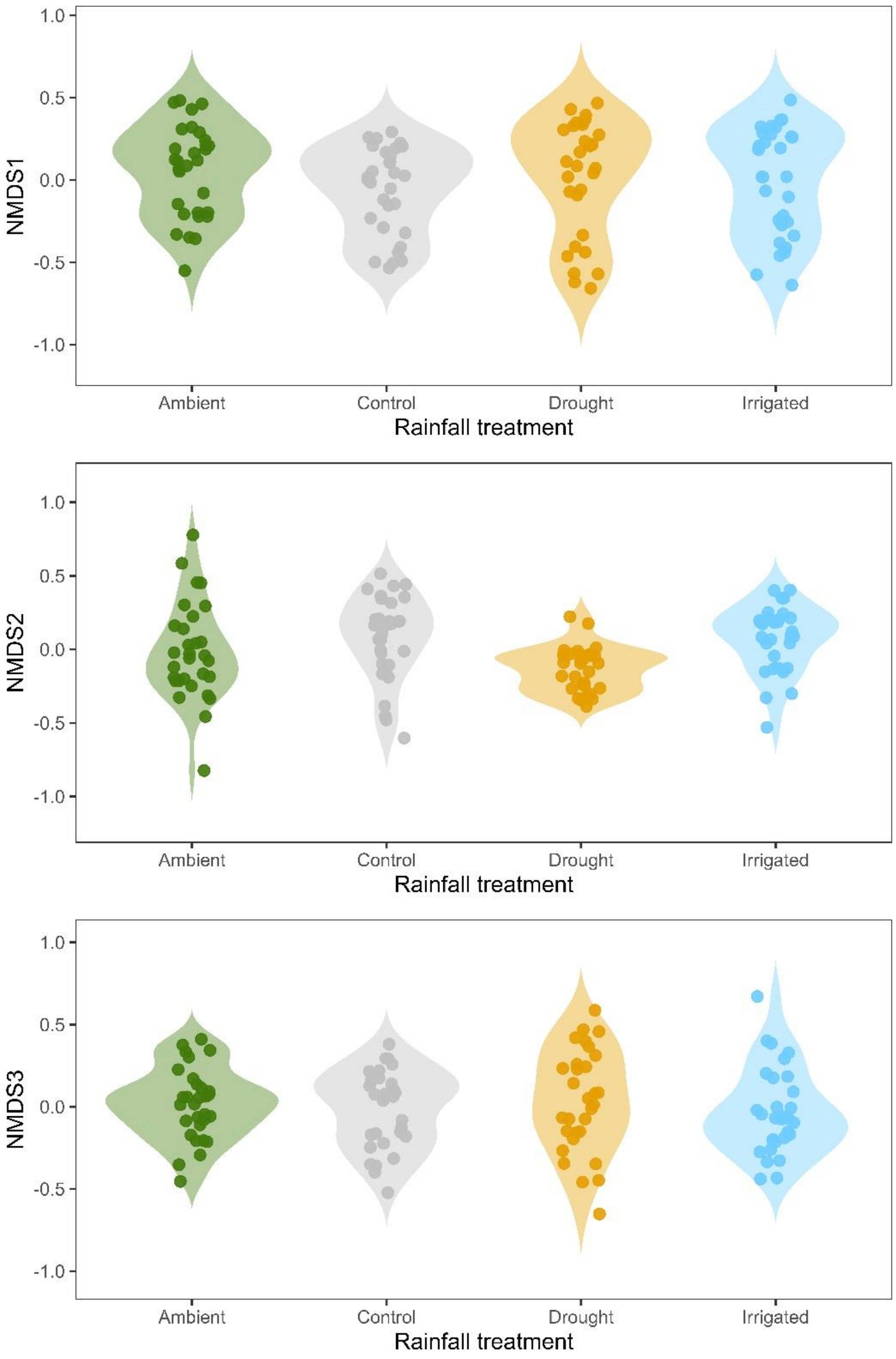


**Figure S5.** No clear difference in community composition between rainfall treatments, as calculated from three axes of non-metric multidimensional scaling.

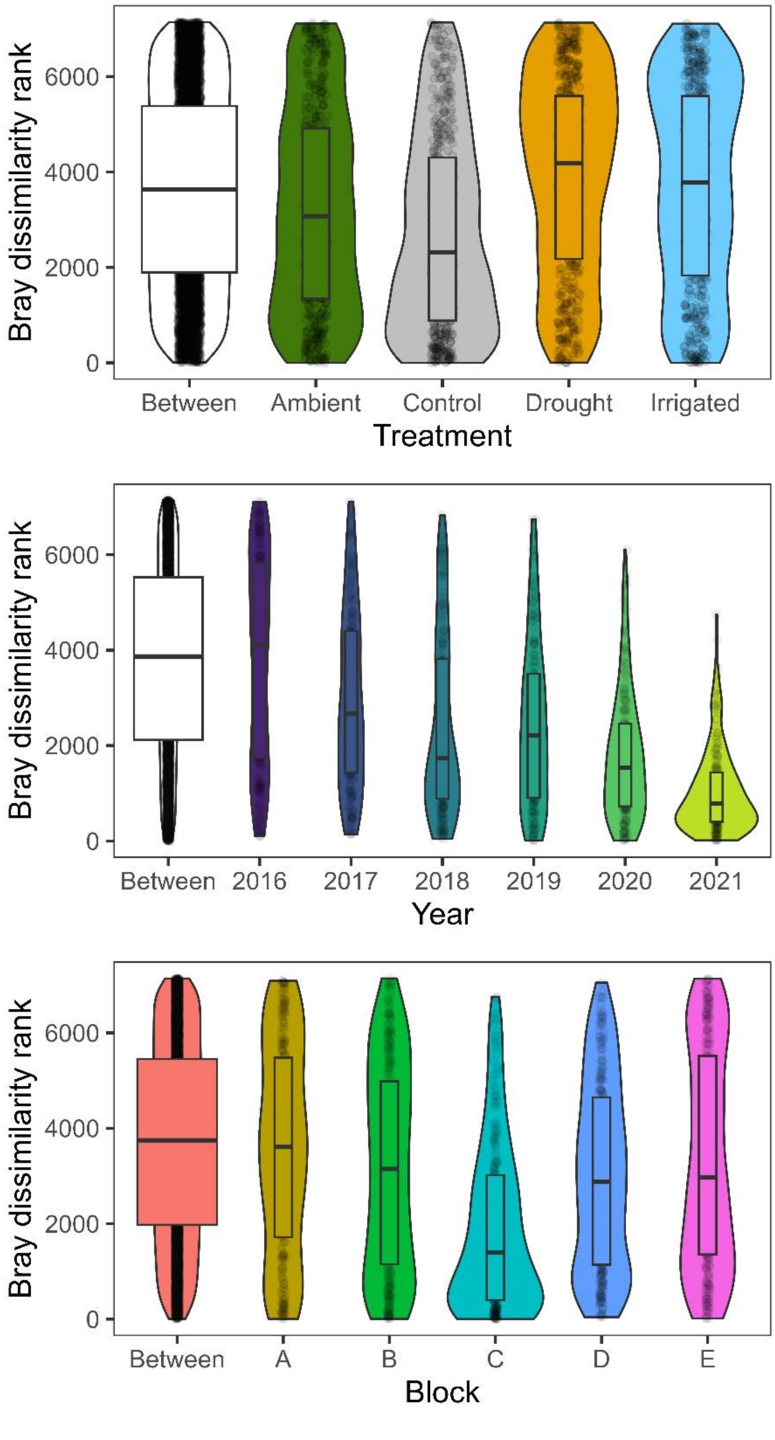


**Figure S6.** Analysis of similarities for within vs, between group community composition comparisons, based on precipitation treatment, observation year, and experimental block.
